# Gene-language models are whole genome representation learners

**DOI:** 10.1101/2024.03.18.585642

**Authors:** Bryan Naidenov, Charles Chen

## Abstract

The language of genetic code embodies a complex grammar and rich syntax of interacting molecular elements. Recent advances in self-supervision and feature learning suggest that statistical learning techniques can identify high-quality quantitative representations from inherent semantic structure. We present a gene-based language model that generates whole-genome vector representations from a population of 16 disease-causing bacterial species by leveraging natural contrastive characteristics between individuals. To achieve this, we developed a set-based learning objective, AB learning, that compares the annotated gene content of two population subsets for use in optimization. Using this foundational objective, we trained a Transformer model to backpropagate information into dense genome vector representations. The resulting bacterial representations, or embeddings, captured important population structure characteristics, like delineations across serotypes and host specificity preferences. Their vector quantities encoded the relevant functional information necessary to achieve state-of-the-art genomic supervised prediction accuracy in 11 out of 12 antibiotic resistance phenotypes.

**Teaser:** Deep transformers capture and encode gene language content to derive versatile latent embeddings of microbial genomes.

## Introduction

The introduction of the self-attention mechanism (*1*), and successively the Transformer architecture, has provided a basis by which neural networks can differentially attend to various parts of the input data, facilitating context-dependent informational exchange. Subsequent Large Language Models (LLMs), like GPT (*2*) and BERT (*3*) have taken advantage of massive unlabeled text corpuses by utilizing self-supervised objectives that explored the natural semantic structure of language. The resulting quantitative word representations, or embeddings, have since become foundational for underlying language representation and enabling high-accuracy text summarization (*4*), document classification (*5*), and sentiment analysis (*6*).

With these successes, new focus has been placed on adapting such models for biological applications by attempting to learn to decode the different sub-cellular languages. In computational protein structure prediction, large successes have been seen with the development of AlphaFold (*7, 8*). Likewise, the prediction of functional properties in proteins has seen significant advancement with the introduction of LLMs (*9*). Genomic applications, however, have seen considerably more limited success due to intrinsic challenges in genomic language that make training more difficult (*10*). Whole-genomes contain ultra-long range language dependencies that cannot fit in standard token-based language models, limiting most deep learning models to predicting functional properties of small base-pair regions (*11-14*). While protein primary structure and protein tertiary structure make natural cross-experimental input-output pairs, contiguous whole-genome sequences lack an established ground-truth label set.

Genes are the fundamental biological units of function and heredity. Their presence, organization, and structure within the genome encode the foundational information to allow an organism to develop, respond to stimuli, and reproduce. Interactions amongst multiple genes, like coordinated gene co-expression (*15*), mimics language in terms of collectively demonstrating higher-order compositional phenomena. Genes intrinsically follow a complicated context-dependent grammar, where the outcome of large gene regulatory cascades (*16*), for example, are dependent on every interaction between the genes involved. Though our understanding of the gene language model has evolved significantly in the last few decades, the comprehensive genome-wide syntax remains enigmatic. The adoption of new language learning technologies, like Transformer models (*1*) and learnable vector-space representations (*3*), provides an opportunity to decipher the meaningful relationships between genes at differing levels of granularity. Specifically, an LLM’s ability to use attention to model interdependent patterns within textual data (*17*), mirrors the continuing need to interpret co-dependence and interactions across genes. As such, LLMs provide an effective statistical learning bridge to interrogate gene feature patterns and use the resulting signals to discover low-dimensional aggregate embeddings of genomes.

We propose a model that adapts self-attention to leverage composite genic information to quantify unlabeled whole-genome sequences. Our methodology, inspired by recent advances in language models, is presented as a deep learning framework capable of exploring gene features in bacterial genome assemblies and deriving meaningful whole-genome quantitative representations that manifest complex biological characteristics. In an exercise to demonstrate its to demonstrate its utility, we target a collection of enteric bacterial species that pose significant public safety risks across several diverse genera including *Salmonella, Escherichia, Vibrio, Campylobacter*, and *Shigella*. We establish that by training the model on the collection of genes already found in the bacterial genome corpus, quantitative embedding can self-organize to form a latent space without relying on corresponding labels. We first introduce our results for embedding a single species, *S. enterica*, before expanding the model’s capabilities in deriving a shared embeddings space for 16 different species. We discuss our deep learning model’s potential to learn from, and encapsulate, the language of gene features as novel quantitative representations capable of elucidating temporal characteristics, characterizing population structure, and inferring important antibiotic resistance phenotypes.

## Results

### Self-organization of the embedding space

To visualize the evolution of genome embeddings over time, we projected the 128-D genome vectors to 2-D vectors using t-distributed Stochastic Neighbor Embedding (t-SNE) (*18*). Prior to training (0 steps), the genome embeddings appeared largely as a point cloud within 2D space (**Fig. 1a**). At 18,000 steps, the embeddings begin to demonstrate signs of self-organization with the embedding space showing stratified genome densities (**Fig. 1b)**. At 24,000 steps (**Fig. 1c**), the embeddings being to form loosely defined clusters; and when the model has converged (160,000 steps, **Fig. 1d**), the clusters became well-defined, and represented an inherent semantic structure of genome embeddings. **Fig. S2a** and **Fig. S2b** illustrate the binary cross entropy loss curve and the positive and negative label accuracies respectively.

**Fig. 1:**
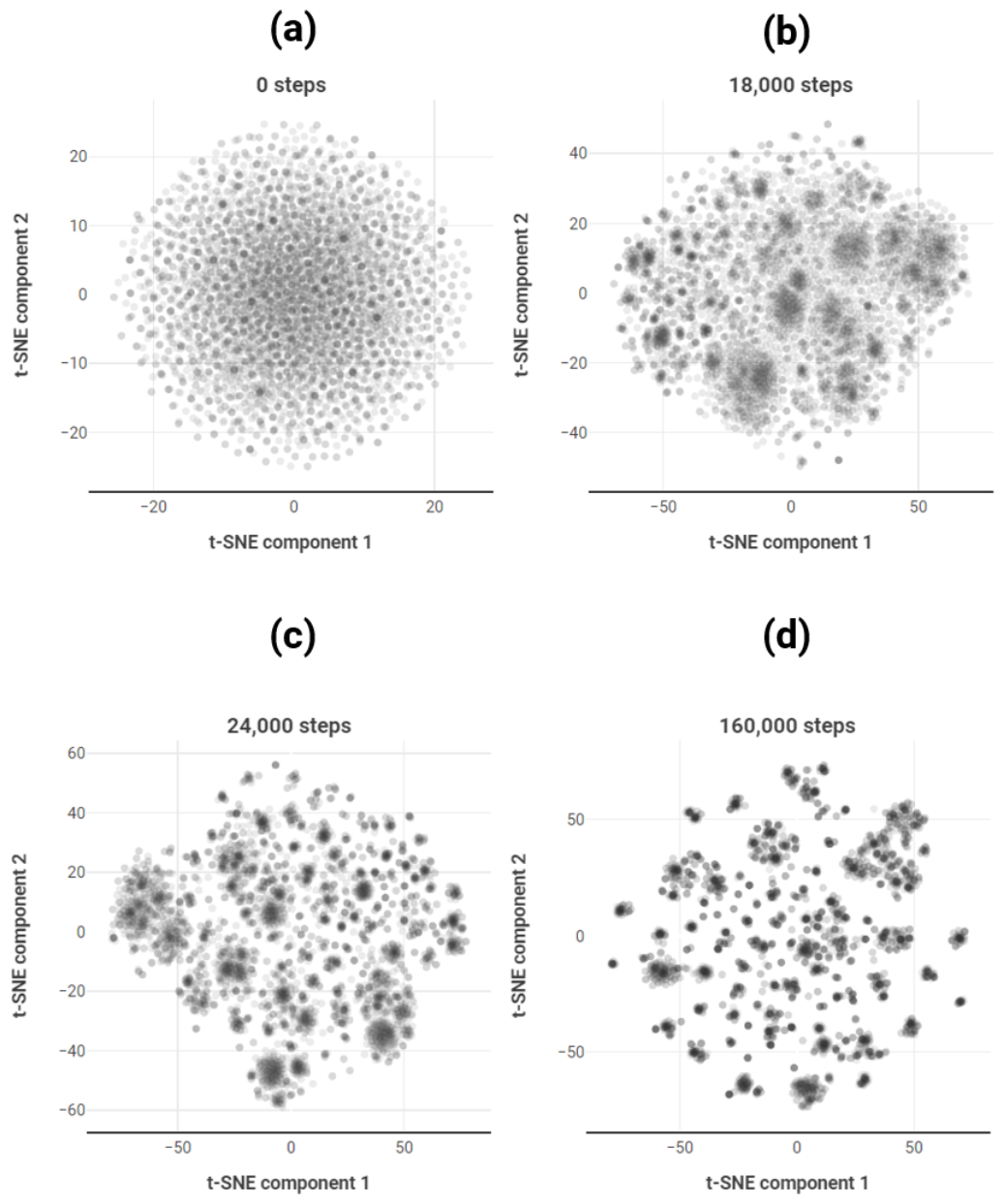
2-dimensional t-SNE representations of 128-dimensional genome embeddings over the course of progressive training steps. Each subfigure shows the genome embeddings projected onto a 2D coordinate plane using t-SNE after training the model for a given number of steps. **a** shows the initialized, untrained embeddings. **b** illustrates the embeddings at 18,000 steps, early in traing. **c** shows the embeddings at 24,000 steps. **d** demonstrates the embeddings after training is complete. As AB genespace training progresses, the embedding representations create a well-stratified embedding space with an inherent semantic structure reflecting the objective function.

### Prediction and characterization of surface antigen structure

To assess the quality of the genome embeddings generated by the Transformer, we interrogated the embedding space for key biological characteristics. Firstly, we examined the distribution of serotype clusters in the learned genomic representations. The complete embedding space, visualized in **Fig. 2a** with color-coded serotypes, exposed a clustering structure of individuals belonging to the same serotype, without having provided any serotype information to the model. The proximity of the various serotype clusters mirrors the Kauffman–White classification groupings. Serotypes clusters composed of individuals found to contain O-antigen groups 6 and 8 (“Hadar”, “Muenchen”, “Newport”, and “Litchfield”, all serovars of the O:8 C_2_-C_3_ serogroup) are seen in close proximity. Likewise, O-6,7,14 antigen-sharing groups like “Mbandaka”, “Braenderup”, and “Infantis” (all members of the O:7 C_1_ serogroup) show up as neighboring clusters in **Fig. 2a**.

**Fig. 2:**
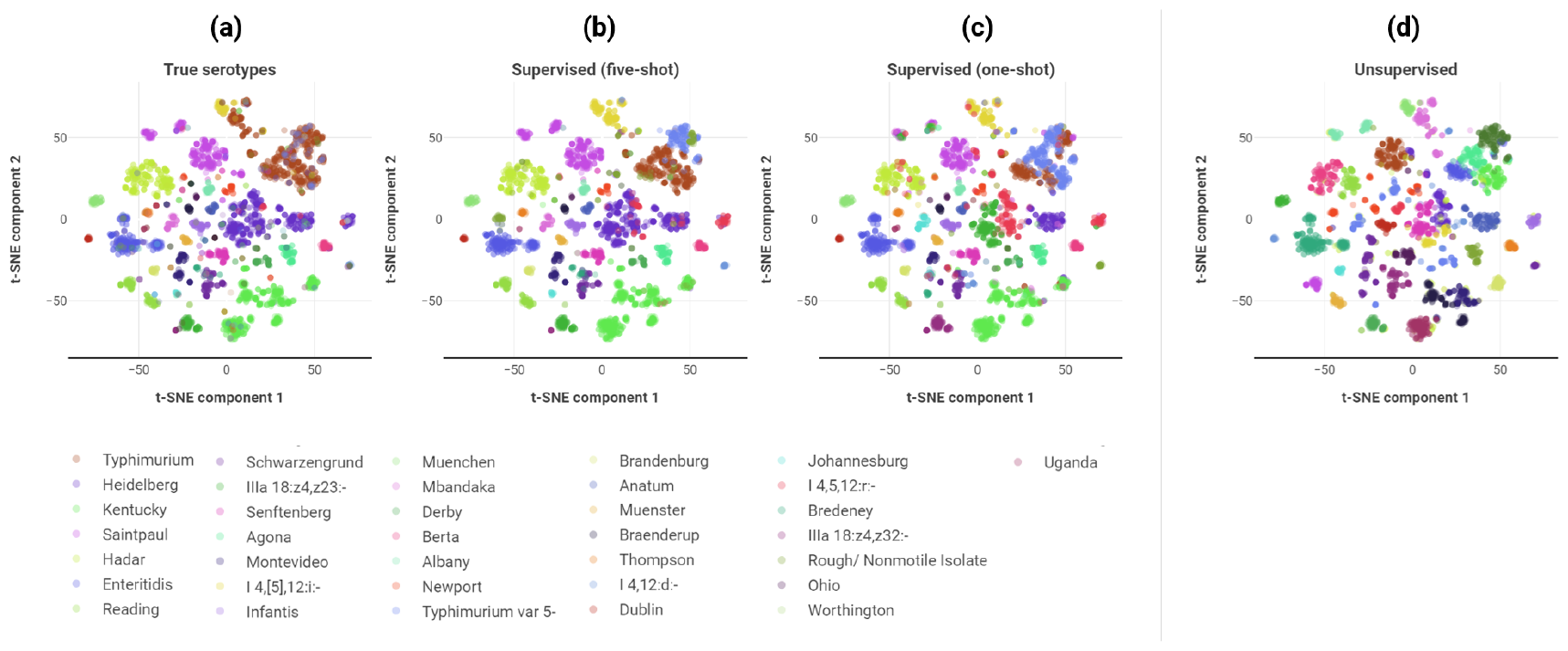
t-SNE of the embedding vectors visualized as 2D points with color-coded serotype information. **a** illustrates the complete serotype space for accessions with serotype metadata. **b** shows one run of the results of predicting serotype labels using few-shot learning (5 representative isolates per serotype in training set). Similarly, **c** shows the results of one run of predicting serotype labels using one representative sample per serotype, i.e. one-shot learning. The legend identifies the major serotype groups. **d** demonstrates the result of using unsupervised clustering to delineate serotypes when no labels are available. **S**ince the clustering method (**d**) has no access to labels, the color scheme does not correspond to the serotype legend.

Using the distinctive serotype semantic structure of the embedding space, we evaluated a supervised model’s ability to directly classify genomes with serotype labels. Using a Support Vector Machine (SVM) with a linear kernel, we trained a model on 50% of each serotype’s samples and achieved an average classification accuracy of 95% (SD=0.24%) on the remaining individuals over 50 randomized replications.

We further demonstrated that embeddings can effectively resolve the serotype space when provided with a small number of representative serotype samples. In a few-shot learning approach (five randomly selected representatives per serotype), the SVM classifier, on average, correctly labeled 79% (SD=4.1%) of the remaining samples over 50 randomized iterations. **Fig. 2b** illustrates one run of resulting serotype classifications overlaid upon the embeddings. Diverse clusters with significant substructure, like “Kentucky”, “Saintpaul” and “Heidelberg”, were reconstructed successfully. It should be noted that a considerable proportion of the error originated from misclassifying the “Typhimurium” serotype as “Typhimurium var. 5”, a highly related Typhimurium variety that is often difficult to differentiate from the standard Typhimurium (*19*).

We further explored low-sample supervision using a one-shot learning paradigm (only one representative individual per serotype). By initializing a K-means (K=number of serotypes) clustering at each representative point, serotype centroids can be identified with no additional information. Then, by sampling the accession closest to the centroid, improved embedding exemplars were acquired. Using these samples, a SVM classifier was trained to correctly label 65% (SD=7.5%) of individuals on average over 50 randomized iterations. **Fig. 2c** depicts a visualization of the outcome of a single round of one-shot learning. Most serotype clusters were accurately reconstructed, including ones with stratification like the “Kentucky” and “Reading” serotypes. Further, most minor serotypes such as “Thompson”, “Uganda”, and “Dublin”) were correctly labeled.

However, due to the limited capabilities of one-shot learning, a greater degree of misclassification can be observed in **Fig. 2c**, primarily due to fragmentation and mislabeling in larger, more diverse serotypes like “Heidelberg”.

Using K-means clustering as an unsupervised approach, we show that major serotypes (limited to serotypes with at least eight members) can be distinguished using genome embeddings without prior examples. In **Fig. 2d**, the unsupervised K-means approach showed that the results aligned with the observed serotype classes from the original agglutination assays. The unsupervised model was able to capture the “Reading”, “Saintpaul”, and “Montevideo” clusters, despite significant sub-clustering. However, larger and more diverse serotypes, like “Kentucky” and “Heidelberg” are overly stratified. The similarity of the class separation produced from the unsupervised embedding method and the true serotypes derived from agglutination assays was measured by the adjusted mutual information metric (AMI). Here, a mean AMI of 0.713 (SD=0.005) was obtained from 200 runs of a randomly initialized clustering model, suggesting the embeddings can be used to identify novel serotypes and also differentiate between distinct serotypes.

### Host specificity, regional localization, and temporal patterns

Preferential host specificity, as it relates to the ability to colonize a particular host species, is a complex, multi-factorial interplay of factors relating to pathogen/host interactions, pathogen virulence (*20*), and host immune system pressures (*21*). **Fig. 3a** showed a well-structured t-SNE visualization that accurately reflects the colonization associated with three meat types: pork chops, chicken breast, and ground turkey. The embeddings were clearly segregated by host preference, with minimal overlap between the dominant poultry clusters. The pork-based accessions form their own smaller clusters and have a minor degree of overlap with both poultry-based clusters.

**Fig. 3:**
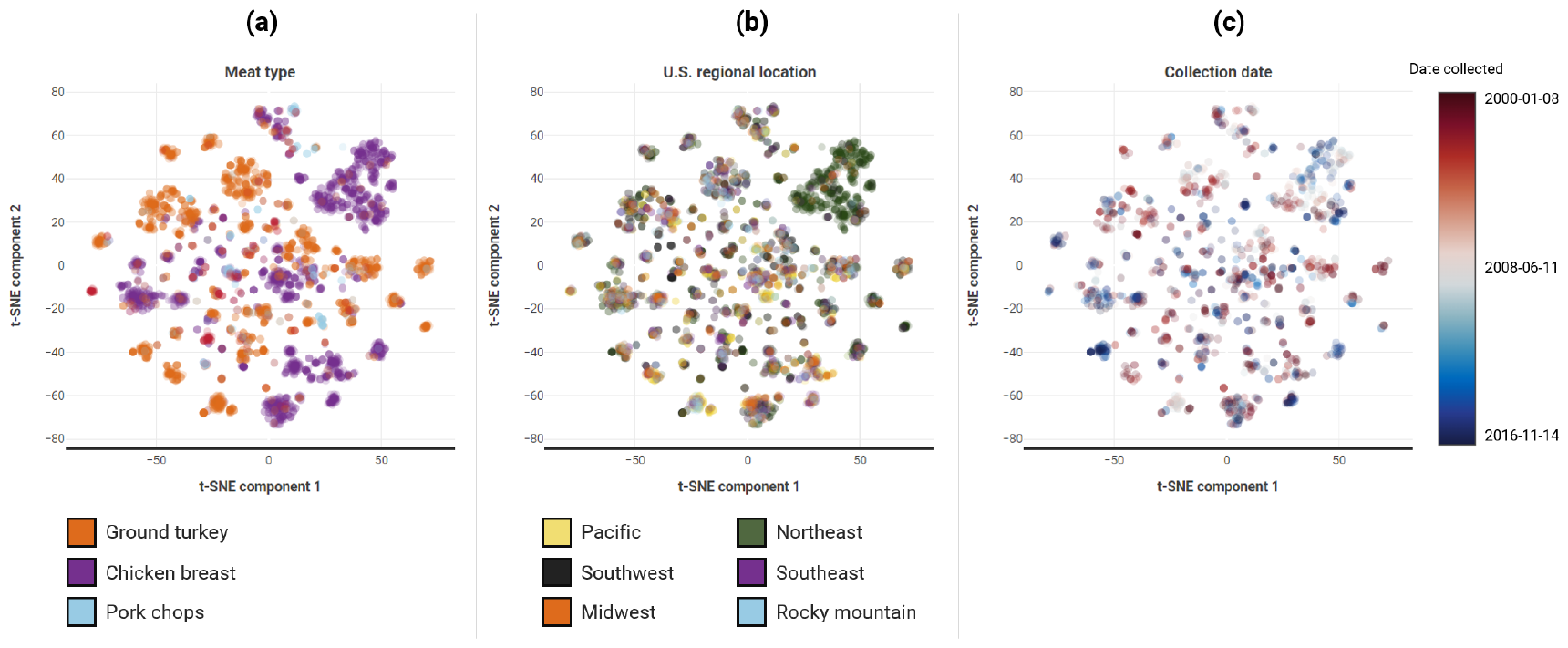
t-SNE representations with points color-coded by key sourcing background. **a** is a t-SNE visualization of the embeddings color-coded by the meat type they were collected from. Here, the three meat type categories are ground turkey, chicken breast, and pork chops. **b** shows a regional color-coding scheme. The color of an accession reflects the U.S. domestic region it was collected from. **c** illustrates the collection data information overlaid upon the embeddings. The red color indicates a collection date closer to 2000, and a blue color indicates that the isolate was collected closer to 2016.

When examining the collection region of each genome (**Fig. 3b**), few obvious clustering patterns were detected, with the exception of a cluster of genomes associated with the northeast American region (Maryland, New York, Connecticut, Pennsylvania). Given the complex production practices, distribution patterns, or handling protocols related to the food supply chain, cluster purity may not be as likely to be observed compared to other attributes.

The collection date information, superimposed over t-SNE projections can be visualized in **Fig. 3c**. Here, the subclustering in the “Reading” serotype was revealed to correspond to a temporal disparity of early 2000 samples and a mid-2010’s outbreak, demonstrating the Transformer model’s ability to capture key temporal genetic changes. Similar observations were made for the “Typhimurium” and “Kentucky” serotypes, with both displaying a subcluster predominantly composed of mid-2010 samples.

### Genomic prediction of antibiotic resistance

Using the embedding vectors as inputs, drug resistance for amoxicillin-clavulanic acid, azithromycin, cefoxitin, ceftiofur, ceftriaxone, ciprofloxacin, gentamicin, kanamycin, nalidixic acid, sulfisoxazole, tetracycline, and trimethoprim-sulfamethoxazole, was predicted. The minimum inhibitory concentration (MIC) phenotypes of 12 common antibiotics were predicted with a logistic regression model as a multi-categorical problem, using 50 randomized iterations of 10-fold cross-validation. **Fig. 4a** illustrated the evaluation of genomic prediction accuracy throughout the AB learning process. It is evident that the prediction accuracy of the supervised model improved as the embeddings were trained, suggesting that our self-supervised learning process provided the embeddings with the ability to classify drug resistance. As shown in **Fig. 4b**, our model significantly improved the prediction capacities for all drugs, except kanamycin, compared to a previously reported k-mer based XGBoost model for the same population (*22*). Our model produced a prediction accuracy of 79.9% compared to 62.4% for the alternative model; and, the embedding-based genomic prediction also yielded significantly narrower 95% confidence intervals.

**Fig. 4:**
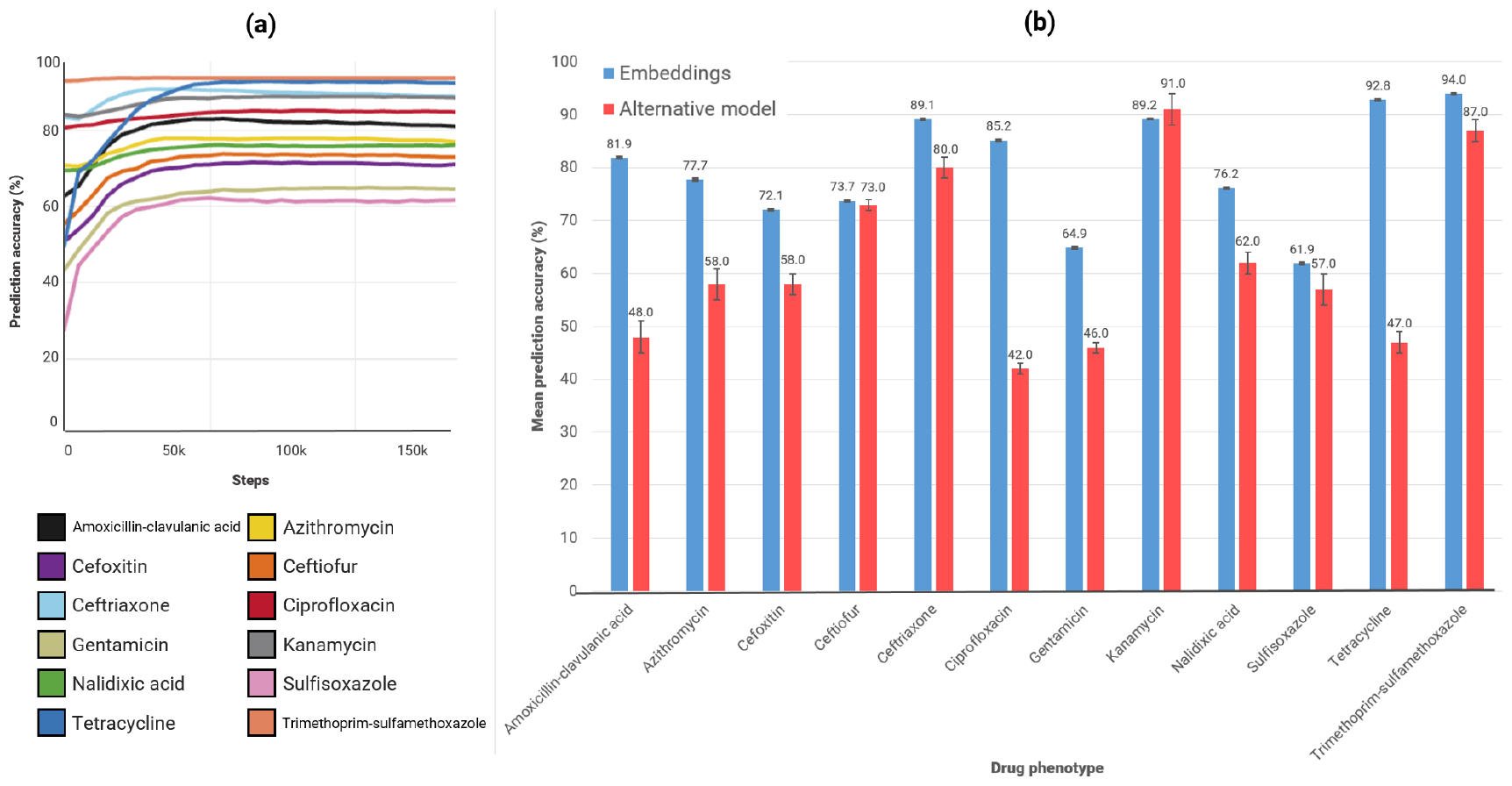
Genomic prediction metrics using embedding vectors as inputs. **a** shows the periodic testing of genomic prediction accuracy over the course of Transformer training. Improvements in predicting the phenotypes corresponding to progress in model training. **b** shows the accuracy results of 50 rounds of 10-fold cross validation. Error bars reflect the 95% confidence interval. Our model (blue) outperforms the alternative model (red) in 11 of 12 phenotypes using the same dataset.

### Taxonomic characteristics of a multi-species embedding space

By expanding the dataset to include genomes from all genera in the dataset, we constructed a multi-species embedding space to examine genetic relationships. **Fig. 5a** demonstrates the capacity of the Transformer model to generate a unified embedding space while preserving taxonomic boundaries at the genus level. Notably, isolates were found to self-organize into distinct clusters at the species level, indicating that embeddings capture subtle genetic differences among closely related bacterial species. Moreover, the *S. enterica* specimens collected from meat tissue, for example, are mostly split from those obtained from the clinical cohort. The trained embeddings were able to summarize variation between species, with respect to the host colony source. Serotype delineations within the species-level clusters are also retained (**Fig. 5b**).

**Fig. 5:**
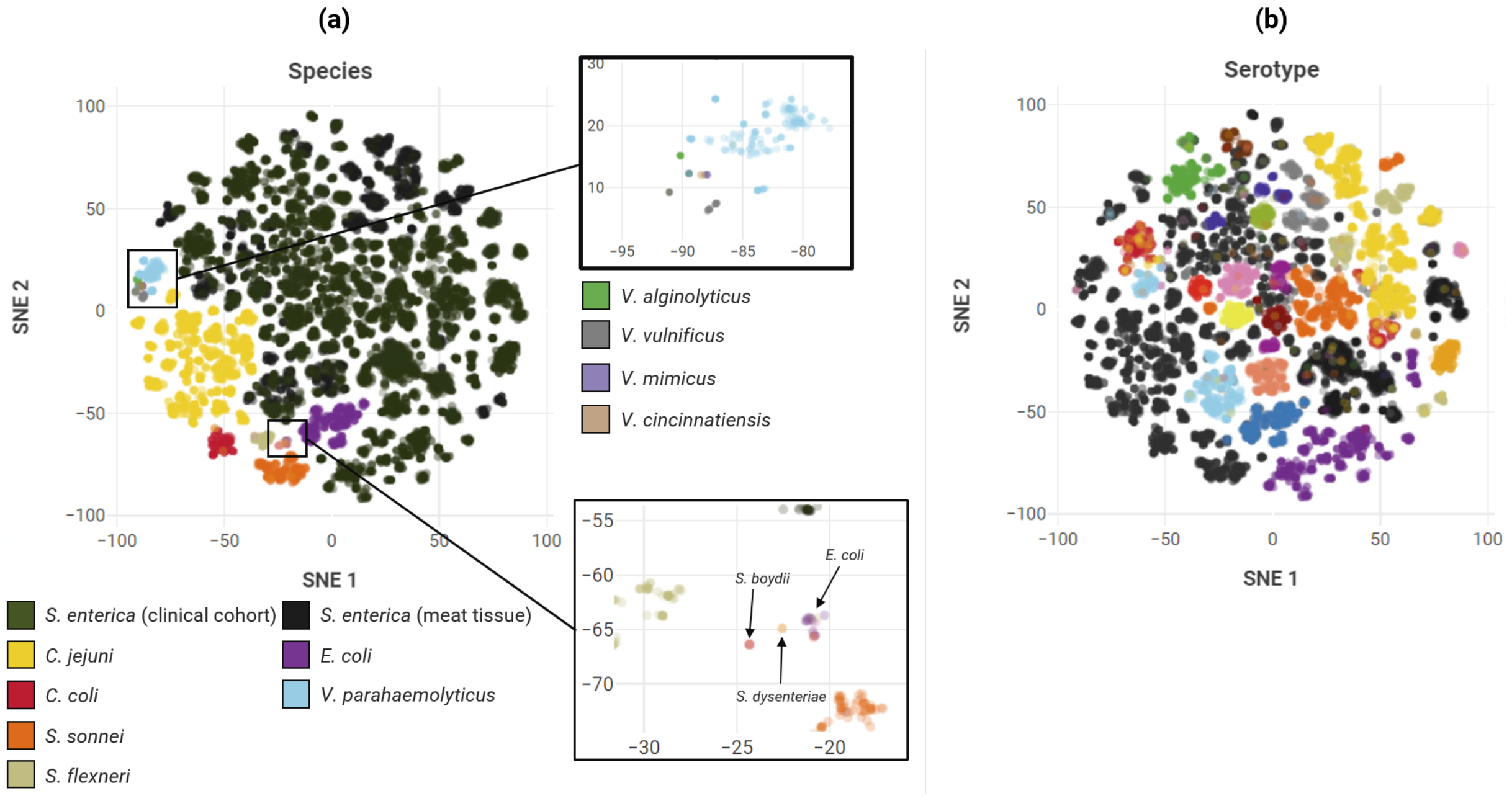
t-SNE of multi-species embedding vectors. **a** visualizes all 16 species (20,145 embeddings) represented by different colors. **b** illustrates the embedding visualizations that known serotype labels.

## Discussion

Much like words in the human language, genomic features represent a rich genetic vocabulary that can be leveraged in comparative classification tasks to propagate implicit supervisory signals. Here we present our AB learning model capable of deriving quantitative encodings of genomes via a self-supervised learning paradigm analogous to those used in recent natural language processing models, like BERT (*3*). Using genomic features as a common genomic vocabulary, and an objective that interrogates the relationship between groups of individuals, we established that composite genome embeddings can be learned without requiring supervised input-output pairs. Moreover, with a large population of assembled and fully annotated genomes as a training “genome corpus”, we demonstrated that an embedding space with meaningful semantic structure can be learned. The resulting whole-genome representations demonstrate a remarkable range of emergent properties, including the ability to distinguish surface antigen configuration, encode discriminative features that accurately predict drug resistance phenotypes, and incorporate multiple distinct genera into a unified embedding space structure.

The self-supervised objective function employed in AB learning framework enables end-to-end training without relying on potentially expensive or time-consuming labeling of genomes, much like how LLMs are able to leverage vast datasets of unlabeled text to train on. We show that our model can translate the gene content of a population to an informative embedding space that can be applied to targeted downstream supervised tasks like predicting drug resistance and serotypes labels. Similar to how prior work has shown that paragraph embeddings aggregate to form topic and category clusters (*23*) based on their word composition, we found that our genome embeddings automatically condense into tight clusters representing important epidemiological properties for the various subsets of the enteric bacteria studied, as seen in **Fig. 2** and **Fig. 3**. The embeddings self-organize into clusters whose proximities are characterized by surface antigen composition and arrangement, in line with the Kauffman–White classification groupings (*24*). In the meat tissue cohort, *S. enterica* serotypes “Bredeney” (antigenic formula 1,4,12,27:l,v:1,7) and “Schwarzengrund” (1,4,12,27:d:1,7) demonstrated an expected close proximity typical of members of the O:4 (B) serogroup (**Fig. 2a**). And, serotypes with O-6,7,14 antigen configurations like “Mbandaka”, Braenderup, and “Infantis” were observed to be in proximity (**Fig. 2a**). Here we also postulate that the embeddings excel at capturing serotype characteristics, even in sample-restrictive scenarios. The observed improvements in classification accuracy when transitioning between one-shot learning (**Fig. 2c**) and five-shot learning (**Fig. 2b**), suggests that an embedding-based classifier can achieve high accuracy with a relatively small training dataset. This ability to perform well with few-shot learning mimics the few-shot characteristics in NLP-based LLMs (*25*).

Furthermore, the embeddings appear to be able to differentiate crucial temporal nuances relating to the collection dates of the samples. For instance, the *S. enterica* Reading serotype is represented by two distinct clusters; isolate dating then confirms that this segregation is entirely temporal in nature; one of the clusters is mostly composed of isolates captured in 2006 – 2008, while the other, larger cluster consists of isolates from 2014 – 2016 (**Fig. 3c**). The model’s partition of the 2014 – 2016 Reading isolates corresponds well with a significant Reading outbreak in 2017 Reading. A time-scaled phylogenetic study pinpointed the emergence of a subclade arising in 2014 – 2016 (*26*), characterized by with distinct genetic profiles, which corresponds closely to the model’s clustering. This distinction is also unsurprising, given that the genetic composition of a population is a function of its temporal and geographic backgrounds (*27, 28*), and parallels can be drawn that word embeddings have been seen to learn to encapsulate historical trends and associated semantic shift in meaning over time (*29*).

Here, we demonstrated that the quantitative elements found within genome embeddings provide state-of-the-art prediction accuracy for MIC phenotypes of 12 drugs using a simple linear model. Prediction of three beta-lactam antibiotics (amoxicillin, cefoxitin, and ceftiofur) yielded consistently high prediction accuracies, ranging between 70 – 82 %, with the fourth beta-lactam being predicted with exceptionally high accuracy (89%) (**Fig. 4b**). These antibiotic resistance phenotypes are characterized by shared beta-lactamase genetic determinates (*30*), like the AmpC (*31, 32*) gene that is present in certain subsets of the population. The results of the prediction performance, and the earlier self-supervised training accuracy, implies that the gene-phenotype relationship was effectively captured and retained in the embedding. The prediction accuracy for two aminoglycosides was mixed. Kanamycin’s MIC values were predicted with a high accuracy (89.2%), while predicting gentamicin poses challenges with a lower accuracy of 64.9% (**Fig. 4b**). Kanamycin’s two dominant MIC categories, 8.0 µg/ml and 64.0 µg/ml, reflected an eight-fold concentration difference and respectively represent a susceptible and resistant phenotype based on breakpoints. In contrast, the two dominate MIC values for gentamicin, 0.5 µg/ml and 0.25 µg/ml, reflect a two-fold increase and both fall within the susceptible phenotype category. The close proximity of these values makes it potentially difficult for the classifier to distinguish between such similar MIC profiles.

Conversely, sulfisoxazole produced the lowest accuracy at 61.9 %. Given that the reported MIC range for sulfisoxazole resides entirely under the susceptible breakpoint (e.g. all isolates were susceptible), the classifier might have struggled to differentiate among these subtle MIC differences. Notably, the remarkable prediction accuracy of ciprofloxacin could be attributed to the fact that its resistance is primarily caused by mutational alterations in target enzymes, specifically substitutions on DNA gyrase genes (*gyrA*) and topoisomerase IV genes (*parC*) (*33*). Since our model does not incorporate any mutational data, its performance seems to suggest that the embeddings might encapsulate certain facets of mutation patterns leading to the resistance, as indicated in prior work (*34*).

In addition to the embeddings characteristics evaluated here, we believe that genome representations could offer additional utilities. Our self-supervised model allows genome representations to materialize by simply providing observations of relevant genetic features, which can easily be identified with annotation software, predicted from upstream models (*11, 35, 36*), or captured by sequencing technologies. The adaptable nature of AB learning as a genomic element comparison, can be extended to multi-omic applications such as transcript abundance and proteomic presence. This expansion permits the generation of novel representations at various levels of the central dogma of molecular biology. These “molecular vocabularies” have the potential to establish unified embeddings or individual embeddings that can subsequently investigated for comparative insights, opening up new avenues for interrogating key biological characteristics and novel properties, beyond epidemiology and medicine.

## Material and Methods

### Population composition and acquisition of bacterial genomic short-reads

Two CDC-sourced bacterial genome repositories were acquired and merged to provide a single large genomic data source for modeling. The first repository was acquired from the NARMS 2017 Retail Meats Surveillance Report (*37*), consisting of 4,585 *S. enterica* BioSample accession numbers collected from contaminated retail meats.

The second repository, containing 15,574 NCBI BioSample accessions from multiple enteric species, was acquired using the “NARMS Now: Human Data” interactive tool (*38*). These samples were collected from a human clinical cohort from 2000 – 2021.

Both CDC repositories were combined into a single bulk paired-end read dataset, totaling 20,159 accession records of 18 species, and including recorded MIC phenotypes for 12 drugs. **Fig. S1** details the complete species breakdown of the consolidated dataset.

#### *De novo* assembly of 20,159 bacterial isolates

SPAdes (*39*), was selected to perform *de novo* whole-genome assembly due to its demonstrated effectiveness at producing highly continuous contigs (*40, 41*) with minimal error rates for microbial genomes, even at low read coverage (*42*). SPAdes was used in combination Trimmomatic with the “—only-assembler” switch. Careful mode (“— careful” switch) was used to improve the assembly accuracy in exchange for incurring additional computational time. See **Table S1** for assembly statistics.

#### *In silco* gene annotation for training data generation

Genes were annotated with Prokka using the included default databases and the Pfam and TIGR HMM databases. AMRFinderPlus was used to identify genes conferring virulence, stress responses, and antibiotic resistance amongst the unresolved ORFs. CD-HIT (*43*) was also applied to unlabeled ORF amino acid sequences, to identify new gene candidate clusters. A sequence identity cutoff of 95% and a minimum 50% length cutoff compared to the representative sequence of the cluster was applied. Clusters with at least 5 individuals were retained. See **Table S2** and **Table S3** for annotation statistics.

### Conceptualization of AB learning as a self-supervised gene-language objective

Here we define a set of *N* genomes (**Equation 1a**), for which each genome contains a subset of all *M* identified genes (**Equation 1b**). For any subset of genomes (**Equation 1c**), the tight genespace (**Equation 1d**) and loose genespace (**Equation 1e**). The resulting genespaces acts as a characterization of the observable genic makeup of the subset.

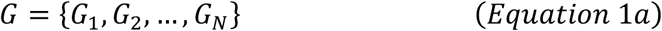

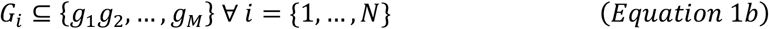

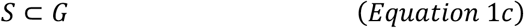

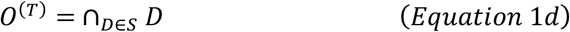

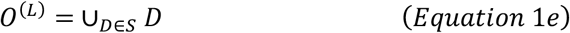

To expose the contrastive characteristics of genome groups, we define the *differential genespace* as the set of genes present in one group’s genespace but absent from another group’s genespace. Given two non-overlapping genome sets, *S*_*-*_ and *S*.(**Equation 2a, Equation 2b**), tight and loose genespaces, 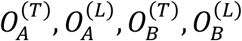, can be computed independently. *Differential tight* and *loose genespaces*, 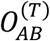 and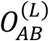, can then be computed by retaining genes unique to the *A*group in each of the genespace specializations (**Equation 2c**). The combined differential genespace specializations form the *AB genespace*. This framework, designated AB learning, provides a self-supervisory signal for learning genomic representations from the population by contrasting genespaces for millions of permuted groupings of individuals to train a Transformer.

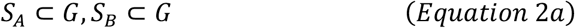

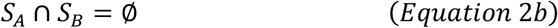

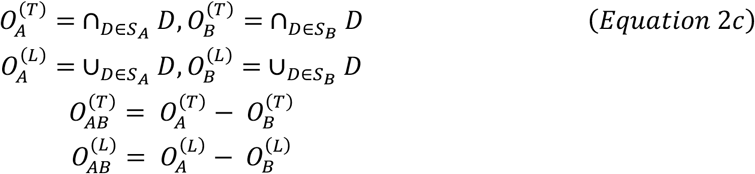

### Transformer architecture and training implementation of a gene language model for embedding genomes

**Fig. 6** provides an overview of the Transformer architecture and input examples. For a single Transformer input sample, accessions are randomly sampled from the population, without replacement, and inserted randomly into two non-overlapping genomes subset pools, *S*_*-*_and *S*.. The total size of either pool is dynamic and can therefore differ between input samples, providing additional permutations of accession groups not possible with fixed pool sizes. Here, *l* is a hyperparameter that species the upper limit of the genome pool sizes.

**Fig. 6.**
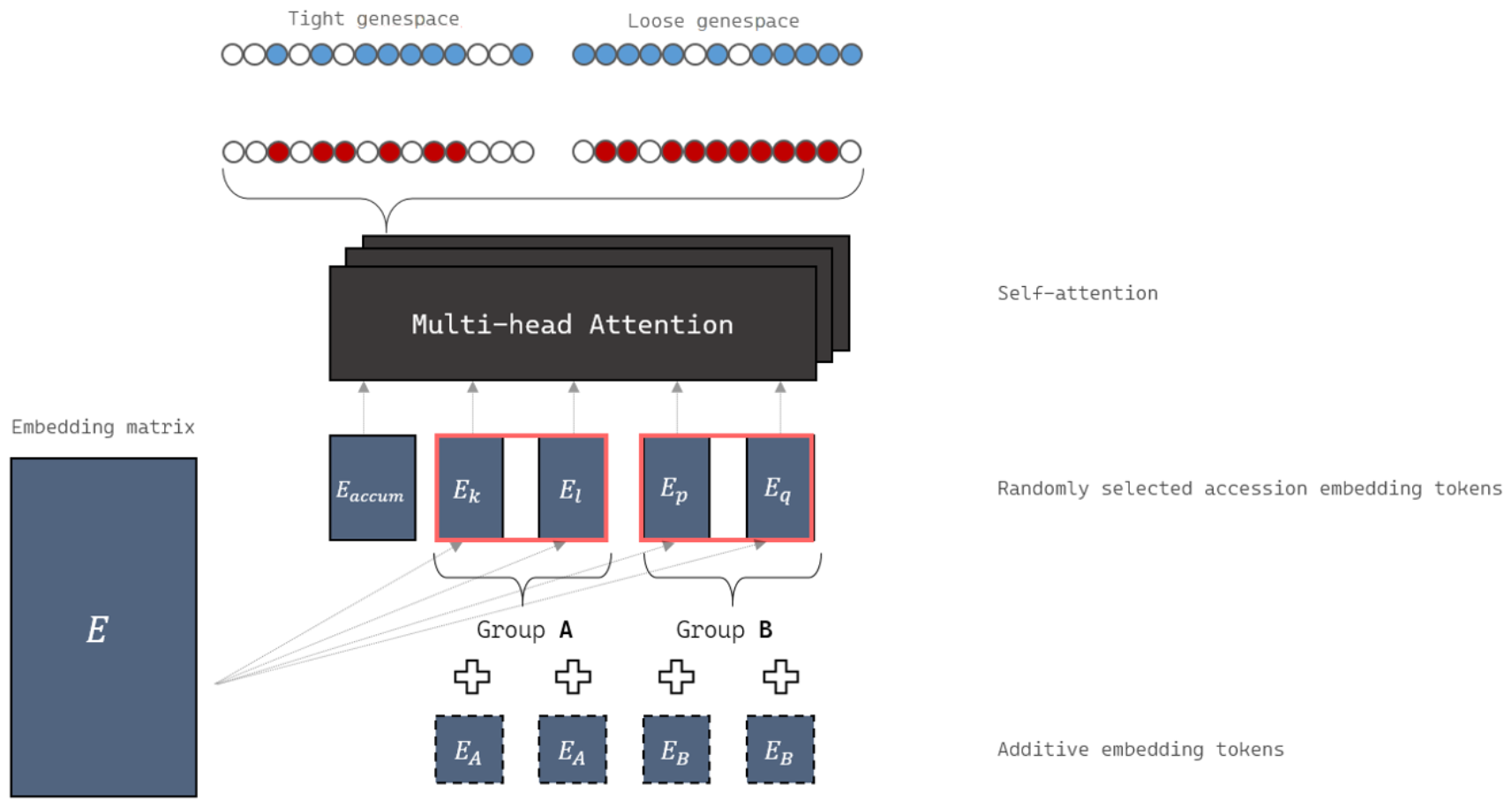
The AB-learning Transformer architecture with 1 = 2. Accession embedding groups are marked with additive tokens (either A or B) and consumed by the Transformer’s attention layer. An accumulator token implicitly captures the relevant genespace information using several layers of attention produce an output AB-genespace that can be compared to the real-world observed AB-genespace.

Genomes are stored as row vectors in a *N*× *d*_*model*_ embedding matrix, where *N*is the number of accessions and *d*_*model*_ the dimensionality of the embedding vectors. Once the *A* and *B* pools are fully populated, the corresponding embedding matrix indices are fetched, and the Transformer input generation stage commences. For each pool, an input index vector of length *l* is populated using the selected embedding indices. When the number of accession indices is less than *l*, the remaining elements of the input index vector are set to a padding token index. Once the two index vectors corresponding to each pool have been populated, the *B* vector is appended to the *A* vector to form a single vector of indices. The rows of the embedding matrix are then indexed by this vector to produce a batch of real-valued accession tokens in a tensor of size *n* × 2*l* × *d*_*model*_, for *n* samples in a batch. An accumulator token is included separately to implicitly aggregate the information necessary to extract and consolidate the *AB* genespace from the other input tokens. The accumulator token has dimensionality *d*_*model*_ and is then prepended to the second axis at the start of the input tensor, producing the final input batch of size *n* × [1 + 2*l*] × *d*_*model*_.

To inform the Transformer as to which tokens belong to the *A*group and which to the *B*, accession tokens are marked by two learnable additive tokens, corresponding to each of the pools.

The body of the Transformer applies multi-head self-attention as described in Vaswani et al. (*1*). Briefly, multi-head attention is a mechanism that uses a set of embeddings to compute the relational structure between inputs allowing for the contextual exchange of information between tokens. With the input embeddings described above (**Equation 3a**), three projection matrices (**Equation 3b**) are used to generate linear projections (**Equation 3c**): query (*Q*), key (*K*), and value (*V*). The degree to which one embedding may “attend” to another embedding (by retrieving a proportion of its projected value vector) can be computed as “attention scores” using the query and key matrices. High similarity between the key and value vectors of two embeddings (taken as the dot-product in one-to-one scenarios), for example, indicates a large degree of information retrieval. In an all-vs-all context, a square matrix of attention scores is computed by matrix multiplication of the *Q* and *K*^**’**^ matrices. The output is the unnormalized attention scores which are scaled by dividing by *d*_4_ (to prevent small gradients) and then normalized to sum to one using the softmax function. The normalized scores are used as weighted sums to determine the influence of each projected embedding in *V*, per-token (**Equation 3d**).

Tokens are transformed by self-attention into context vectors that represents the weighted sum of other embedding value projections. This process is performed *H* attention heads allowing each head to specialize in a particular part of relational information retrieval (**Equation 3e**). For performance reasons, the attention heads operate in a reduced dimensionality and concatenate the transformed tokens such that the dimensionality of the output context embeddings is equal to the input embedding dimensionality. The tokens are then processed by a 2-layer multilayer perceptron with GELU activations (*44*) prior to the next attention layer.

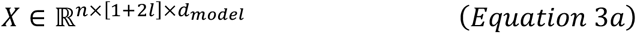

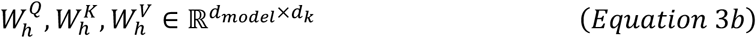

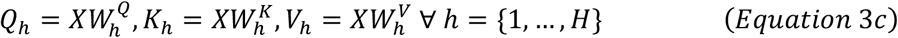

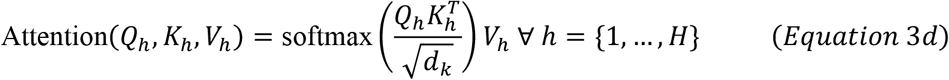

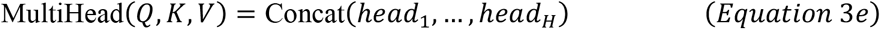

AB genespace outputs are generated by using the accumulator token as an input to a binary classification layer. The calculation of the binary cross entropy loss function over the batch is described in **Equation 4**. Given a batch of *n* samples, the predicted *AB* genespaces (produced from the linear classification layer of the accumulator tokens) are compared with the observed *AB* genespaces. Here, Ŷ is a *n* × 2*M* matrix representing the predicted probabilities for the AB genespace from the model inputs. *Y* is a *n* × (2*M*) binary matrix where indices pointing to genes known to be present in the *AB* genespace are 1 and genes absent from the *AB* genespace are 0.

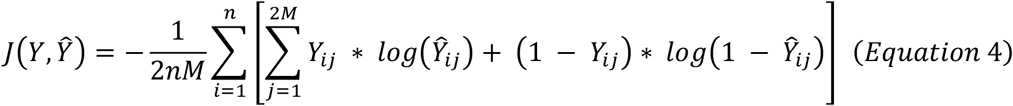

### Optimization, learning rate scheduling strategy, hyperparameters, and software utilities

For parameter optimization, AdamW (*45*) was used with β_1_ = 0.9, β_2_ = 0.999, a weight decay of 0.2, and a projection dropout probability of 10%. As with other language models (*3, 46, 47*), a learning rate scheduler was applied to gradually increase the learning rate for an initial 5,000 batches (steps) before decreasing it towards zero as the training completes. A batch size of 2,048 samples was used, as large batches improve generalizability in self-supervised tasks (*3, 25, 48, 49*).

The upper limit on genome pool size, *l*, was set to 5. An embedding dimensionality of *d*_*model*_ = 128 was used with 4 attention heads and 4 attention layers. Default PyTorch weight initializations were used for embedding and function parameters. The model construction and training were performed using the PyTorch (*50*) deep learning library. Four V100 Volta NVIDIA GPUs were used to train the model for 165,000 steps with FT16 mixed precision.

Visualizations were constructed with Plotly (v5.14.1) (*51*). NumPy (*52*) (v1.21.5) and SciPy (*53*) (v1.7.3) were used for data transformation and manipulation. Pandas (*54*) (v1.2.3) was utilized as a tabular interface. The Scikit-learn (*55*) (v1.0.2) Python package was used to apply machining learning and statistical models to harvested embeddings.

Biopython (*56*) (v1.79) was employed for FASTA sequence manipulation.

## Author contributions

Conceptualization: BN, CC

Methodology: BN, CC

Data processing and bioinformatics analysis: BN

AB learning code development: BN

Visualization: BN

Supervision: CC

Writing: BN, CC

## Data and material availability

The metadata file, phenotypes of the minimum inhibitory concentration (MIC) and a Zarr file containing the differential AB genespace used in this study can be accessed via Dryad link-https://doi.org/doi:10.5061/dryad.vx0k6djzn. And, AB learning algorithm is available at https://github.com/bryan-n-arch/AB-learning.

