## Supplementary material for "Gene-language models are whole genome representation learners"

**This file includes:**

Section S1: Isolate name resolution and procurement of reads from Sequence Read Archive
Section S2: Read trimming and de novo whole-genome assembly with SPAdes
Section S3: Alignment to the UniPort Knowledge Base to identify known prokaryotic genes
Section S4: Integration of the Pfam and TIGR high-quality hidden Markov model (HMM) databases to detect conserved homology
Section S5: AMR-specific gene annotation by scanning against NCBI reference gene collection
Section S6: Population-wide ORF clustering to consolidate amino acid sequences with high similarity
Section S7: Multi-headed attention layer and attention masking
Section S8: Transformer output layer
Fig. S1: The species breakdown for the complete bacterial dataset.
Section S9: De novo bacterial genome assemblies
Table S1: Major species group genome assembly statistics.
Section S10: Gene annotation and pan-genomic characteristics
Table S2: Major species group gene annotation statistics
Table S3: Core-gene, shell-gene, and cloud gene counts for each major species group
Fig. S2: Training statistics over the course of on S. enterica as a single-species model

### Section S1: Isolate name resolution and procurement of reads from Sequence Read Archive

For all accessions provided by NARMS, corresponding BioSample identifiers were resolved to NCBI Sequence Read Archive (SRA) identifiers. For this, NCBI servers were queried in a two-step process using the esearch (v16.5) and efetch (v16.5) utilities from the Entrez Direct (EDirect) Unix software suite(*1*). For each accession, esearch was used with the "–db sra" argument and the "-query" switch followed by the BioSample string with the keyword "[biosample]" appended. The resulting server response was then piped into the efetch software with the "–format runinfo" parameter so as to extract the downloadable "SRA Run".

After name resolution, short-read sequence data was downloaded using the NCBI SRA Toolkit (v2.10.8) software suite(*2*). The included prefetch utility was used to request each SRA Run by passing its accession as an argument. In total, 20,159 SRA requests were submitted and successfully downloaded from NCBI. To convert the compressed SRA binary files to a consumable FASTQ format for downstream trimming and assembly, the fasterq-dump utility, from the Toolkit, was used with the SRA files as the input argument. The resulting files were unpacked as pairs of paired-end Illumina reads in FASTQ format.

### Section S2: Read trimming and de novo whole-genome assembly with SPAdes

Read trimming was performed following the recommended procedure established by the SPAdes developers to yield improved *de novo* assembly results. As illustrated in several SPAdes benchmarks, deactivating the in-built SPAdes read correction and adjusting several parameters can produce improved whole-genome assembly results(*3*). For this, Trimmomatic(*4*) was to perform both adapter removal and base-pair trimming. For the removal of Illumina-specific adapter sequences, Trimmomatic (v0.39) was executed with the "ILLUMINACLIP" augment specified with a seed mismatch of 2, a palindrome clip threshold of 30, and a simple clip threshold of 10. For read trimming, the "SLIDINGWINDOW" argument with a window size of 4 and a required quality minimum of 20 was applied. Finally, the leading and trailing sections of each read were further trimmed using the "LEADING" and "TRAILING" arguments with a quality specification of 3. From the input paired-end reads, Trimmomatic produced the trimmed left and right pairs along with trimmed left and right unpaired reads (reads that could not find a paired mate due to either mate being filtered out).

The de Bruijn graph-based prokaryotic assembler, SPAdes(*5*), was selected to perform *de novo* whole-genome assembly due to its demonstrated effectiveness at producing highly continuous contigs(*6, 7*) with minimal error rates for microbial genomes, even at low read coverage(*8*). With the reads already trimmed, SPAdes' (Version 3.1.5.3, 64-bit prebuilt Linux binary) internal read-correction procedure was selectively deactivated by applying the "—only-assembler" switch, as suggested by the developers. SPAdes was executed in careful mode ("—careful" switch) to improve the assembly accuracy in exchange for incurring additional computational time. To maximize data efficacy, we utilized SPAdes' capabilities in assembling single-unpaired reads emitted by Trimmomatic; we specified the trimmed paired-end reads as options -1 and -2 (for left and right, respectively) and two usages of the "–s" switch for each single unpaired read files emitted by Trimmomatic.

<u>Identification of open reading frames for protein-coding genes</u>

To characterize the gene content in the bacterial assemblies, a multi-step gene annotation pipeline was established. Prokka(*9*) is a bacterial annotation software that utilizes a suite of gene feature prediction tools for marking proteins coding genes and providing functional information. We developed an extended Prokka-based (v1.14.5) annotation pipeline with several augmentations to improve gene detection.

As a first step, Prokka can identify the base-pair coordinates for predicted open-reading frames independently across each genome assembly. For this, the gene-finding software Prodigal(*10*), was used to mark likely candidate genes for downstream functional annotation by scoring potential open reading frames. Open reading frames (ORFs) that passed Prodigal's scoring system were extended until a stop codon was reached providing the complete nucleotide range for each hypothetical gene. The contig localization and base-pair coordinates for these hypothetical genes were retained to provide context for additional annotation steps later in the pipeline. Prokka command line switches were left to the default options. A Singularity(*11*) (v2.5.2) shell was used to interact with Prokka to facilitate a permissible Linux environment for the installation of additional databases later in the procedure.

**Section S3: Alignment to the UniPort Knowledge Base to identify known prokaryotic genes**

For all marked hypothetical genes, the corresponding translated amino acid sequence was matched against the bundled UniProt(*12*) database for genes that have been with real protein or transcript evidence. This operation was performed internally within the Prokka annotation pipeline using BLAST+(*13*) (v2.9) to search against the UniProt "sprot" database (Release 2019_09) for all proteins contained within the bacterial kingdom. For an amino acid similarity hit to be significant, and thus reported as a successful match, a default Prokka e-value of $1 \times 10^{-6}$ was used as a threshold criterion. Prokka, by default, also scans the genome for insertion sequences using ISFinder(*14*) database and for antibiotic genes using Antimicrobial Resistance Reference Gene Database (BARRGD). Non-coding DNA was not considered.

**Section S4: Integration of the Pfam and TIGR high-quality hidden Markov model (HMM) databases to detect conserved homology**

In addition to using the UniProt database for searching for matches based on amino acid similarity via BLAST-based alignment, the Prokka pipeline provides an option to use a more sensitive hidden Markov model(*15, 16*) (HMM) for identifying matches to protein domains from the ORFs that had no significant hits in the UniProt database. In biological sequences, the profile hidden Markov model is a probabilistic model that is used to statistically model the transition probabilities between matched/unmatched nucleotides, insertions, and deletion gaps derived from prior multiple sequence alignments. Characteristically, this type of gene annotation methodology is significantly more robust to sequence gaps and more grounded in evolutionary divergences when compared to conventional alignment-based methods like BLAST(*17*). To search for the presence of homology to known conserved protein domains from the ORFs that did not map to UniProt genes, Prokka employs the "hmmscan" command from the HMMER3(*18*) software distribution. hmmscan will query the translated protein sequences against collections of protein profiles within any HMM profile database. A single database may contain any number of protein profiles; as such, a protein match will only be significant if the HMM produces an e-value of less than or equal to 1e-06(*9*).

In Prokka, the HMM step is only applied with a single HMM profile database, the High-quality Automated and Manual Annotation of Proteins(*19*) (HAMAP) database. Since the HAMAP database is limited to just 2,388 protein family profiles (https://hamap.expasy.org/, UniProt release 2023_01, 22-Feb-2023), its singular usage would likely miss many potential protein targets. To improve the efficacy of the detection capabilities of Prokka, two supplementary HMM peptide databases (Pfam, TIGRFAM) were integrated into the Prokka annotation kit, in addition to HAMAP. The Pfams-A(*20*) database, containing 19,632 protein families, was selected given its high-quality peptide profiles for domains and protein families that have been manually catalogued by experts. HMM profiles (v35.0 November 2021 release) were acquired via the official FTP server (http://ftp.ebi.ac.uk/pub/databases/Pfam/current_release/Pfam-A.hmm.gz). A second database, TIGR (Version 7.0) profiles was also acquired from the NCBI FTP server ((https://ftp.ncbi.nlm.nih.gov/hmm/7.0/hmm_PGAP.HMM.tgz), representing 15,346 protein families. Integration of these two database was performed using the Prokka "–setupdb" switch, by applying the "hmmpress" indexing command from HMMER3.

**Section S5: AMR-specific gene annotation by scanning against NCBI reference gene collection**

To more comprehensively identify putative genetic drivers of drug resistance, an additional step was taken to annotate the AMR-specific gene content of each microbe. AMRFinderPlus(*21*) was used to identify genes conferring virulence, stress responses, and antibiotic resistance amongst the unresolved ORFs. Such genes might be missing within the general-purpose UniProt Knowledge Base and the HMM profiles described earlier. Internally, AMRFinderPlus uses a mixture of manually specified BLAST cutoffs (BlastRules) and carefully selected HMM AMR profiles.

In order to avoid overwriting genes that had been successfully annotated from the earlier steps, the gene detection phase of AMRFinderPlus was applied exclusively to unannotated ORFs previously identified by Prodigal. For each accession, the nucleotide sequence ranges from the unmatched ORFs were extracted and deposited into a per-accession FASTA file to accommodate AMRFinderPlus's input interface. AMRFinderPlus was executed using the "—plus" switch to incorporate additional prokaryotic database entries and the "–n" switch to specify nucleotide-level annotation. From the output table, significant AMR hits were used to update the original GFF3 output from Prokka, replacing hypothetical genes designations with named AMRFinderPlus features.

**Section S6: Population-wide ORF clustering to consolidate amino acid sequences with high similarity**

Even with the above steps, a subset of ORFs were expected to remain unlabeled as the various protein knowledge databases cannot capture the complete space of naturally existing genes. By leveraging the prior high-confidence ORF predictions from Prodigal, putative genes can be retained by identifying ORFs with some degree of redundancy within the population. In an effort to retain potentially significant gene features, these unlabeled ORFs were clustered using amino acid similarity. To achieve this, the nucleotide sequences from the unannotated ORFs were extracted, based on the coordinates provided by Prodigal. Their base-pair representations were then translated into amino acid (AA) sequences with the reverse complement transformation applied depending on the strand parameter emitted by Prodigal. The resulting AA sequences were deposited into a FASTA file for bulk clustering.

For high-confidence ORFs, if a significant level of population-wide redundancy is identified (as indicated by the size of cluster membership), said ORF can be reclassified as a putative gene and retained in the dataset. CD-HIT(*22*) (Cluster Database at High Identity with Tolerance) is a high-throughput clustering software designed for comparing biological sequences. By leveraging a short word filter matching with greedy incremental clustering, haCD-HIT can efficiently produce a set of sequence clusters from a specified sequence file subject to user-defined threshold criteria. The "cd-hit" utility, within the CD-HIT suite (v4.8.1), was applied to the FASTA containing the unlabeled ORF amino acid sequences. The "–c" switch was used as with a value of 0.95 to define a user sequence identity cutoff of 95% while the "–s" switch was used with a 0.50 value to define a minimum 50% length cutoff compared to the representative sequence of the cluster. The "–g" option was used with a 1 argument to maximize accuracy. The resulting CLSTR file was parsed to identify ORF clusters that met the above criteria and were found in at least 5 individuals. In the annotation file, formally unannotated ORFs were collectively renamed based on their cluster membership.

**Section S7: Multi-headed attention layer and attention masking**

Attention masking is used to prevent padding tokens from interacting with self-attention. Without masking, padding tokens create responses in the unnormalized scores calculations, which will attract some degree of attention from non-padding tokens. For this, the padding tokens' unnormalized attention scores are set to $-1 \times 10^4$ prior to the softmax operation during the forward pass. This produces post-softmax values close to zero for padding tokens (lowest stable value for 16-bit floats), ensuring that they are not considered in the attention layers.

**Section S8: Transformer output layer**

After applying several layers of attention, the model is then required to reconstruct the $AB$ genespace vector for each input sample in the batch. The transformed accumulator token is used as input to a linear classification layer to produce unnormalized class logits. Class scores can then be generated by applying a sigmoid function to the unnormalized logits, limiting the output to a range between 0 and 1. The scores represent the predicted probabilities for the presence of each possible gene in the $AB$ genespace. In this case, the output for a given input token batch will be of shape $n \times (2M)$ representing the $AB$ genespace containing $M$ differential tight genespace genes and $M$ differential loose genespace genes.

For the observed $AB$ genespace, the 4 total genespaces (one tight and loose for each pool) are used to generate 2 differential genespaces with respect to the input accessions. We vectorized and concatenated the tight and loose differential genespaces to form a single $AB$ *genespace*. The difference between the model's $AB$ genespace and the observable $AB$ genespace provides a foundation with which to compute loss for weight optimization.

Binary cross entropy was used to compute the aggregate batch loss by comparing the distance between the predicted $AB$ genespace and the observed $AB$ genespace. In this case, the presence of genes in the $AB$ genespace represent multiple independent categories that can be compared to the normalized probabilities emitted from the classification layer with the input accumulator token.

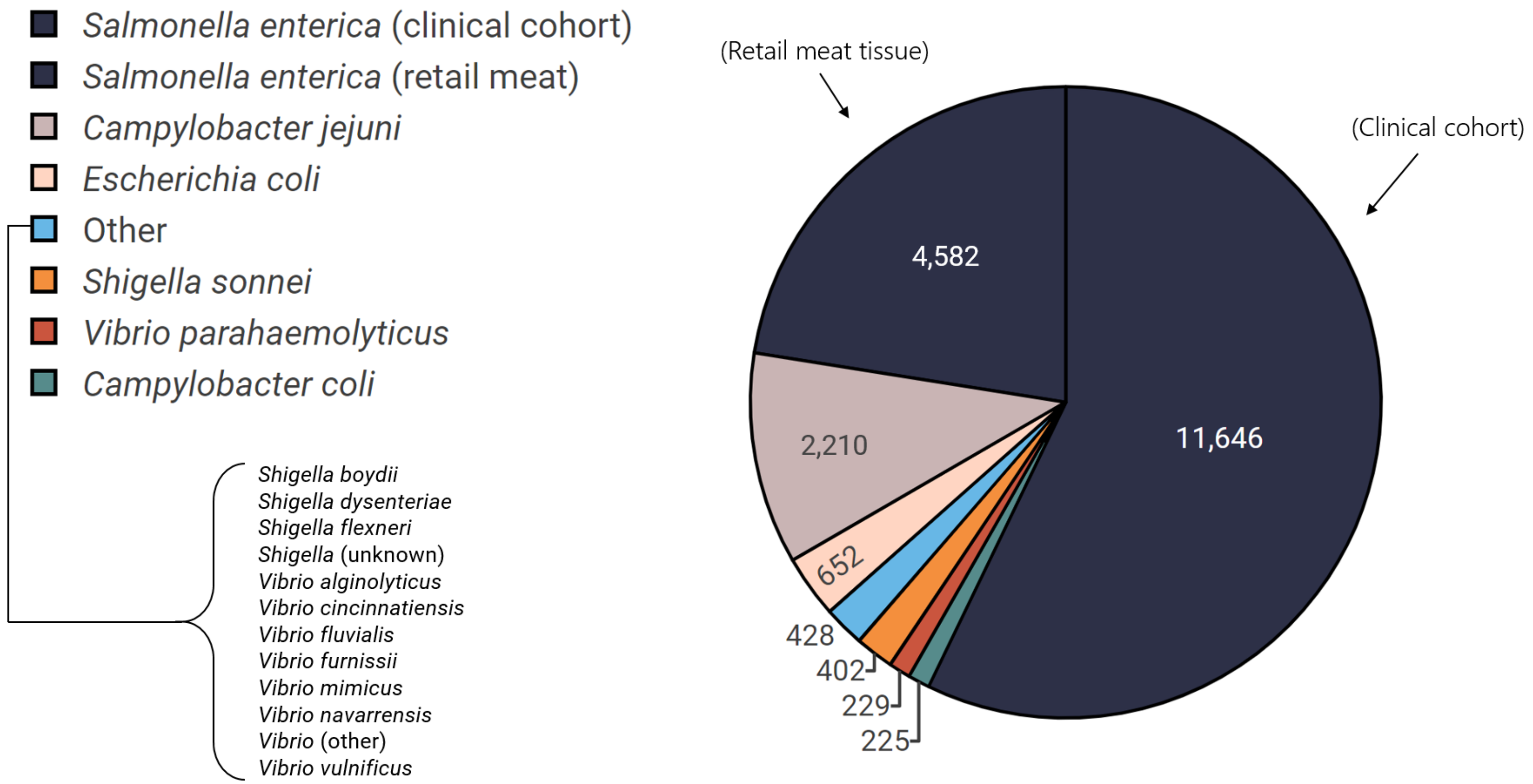


**Fig. S1**: **The species breakdown for the complete bacterial dataset.** There are 16 species in total. Species with few samples are grouped into the "other" category (light blue).

### Section S9: De novo bacterial genome assemblies

**Table S1** summarizes quality metrics that describe the completeness and accuracy of the assembled accessions. Both *S. enterica* cohorts produced comparable assembly summary statistics, with a mean genome size of 4.9 megabases (MB) being within the range of other reported *S. enterica* genome sizes[15, 71, 72]. Similar results were seen for *V. parahaemolyticus* (5.2 MB)(*23, 24*) and *S. sonnei* (5.0 MB)(*25*). The *E. coli* assemblies produced the largest average genome size (5.7 MB), while the sizes of the *Campylobacter* species, *C. jejuni and C. coli*, were the smallest; both matched prior reported size values(*26*). The GC content also matched the reported standards; our results indicate that the *S. enterica*, *E. coli*, *S. sonnei*, *V. parahaemolyticus*, *C. jejuni*, and *C. coli* assembly groups match the reported mean GC content statistics of 52%(*27-29*), 50%(*30, 31*), 51%(*32, 33*), 45%(*34, 35*), 30%(*36, 37*), and 31%(*36, 38*), respectively. As seen in **Table S1**, the lowest N50 is 24,478 (*S. sonnei*), which is significantly higher than the average prokaryotic gene length of approximately 924 bp(*39*), suggesting these assemblies are suitable for annotation.

| Average (SD) | *S. enterica* meat tissue | *S. enterica* clinical cohort | *E. coli* | *S. sonnei* | *V. parahaemolyticus* | *C. jejuni* | *C. coli* |
|---|---|---|---|---|---|---|---|
| Genome size | 4,907,797 (240,060) | 4,860,290 (424,078) | 5,668,713 (489,696) | 4,984,984 (422,814) | 5,192,820 (395,058) | 1,718,240 (119,731) | 1,793,211 (214,774) |
| N50 | 313,224 (61,798) | 399,613 (193,445) | 155,852 (45,554) | 24,478 (2,978) | 504,144 (261,179) | 169,778 (121,316) | 228,476 (176,444) |
| Contig count | 178 (292) | 229 (737) | 810 (711) | 1074 (644) | 172 (340) | 93 (512) | 168 (304) |
| GC content | 52.1% (0.6%) | 52.1% (0.2%) | 50.4% (0.2%) | 50.8% (0.1%) | 45.3% (0.3%) | 30.5% (0.4%) | 31.3% (0.4%) |

**Table S1: Major species group genome assembly statistics.** Listed are the means and standard deviations of the assembly size, N50, contig counts, and GC (guanine/cytosine) content.

### Section S10: Gene annotation and pan-genomic characteristics

A total of 33,091 distinct genes were identified across all analyzed species. The average annotated gene counts were similar across *S. enterica*, *E. coli*, *S. sonnei*, and *V. parahaemolyticus*, resting in the range of 3,400 – 4,300 (**Table S2**). The two *Campylobacter* species, *C. jejuni* and *C. coli*, produced significantly lower mean gene counts (1,400 – 1,500), reflective of their smaller genome sizes. For all species, the UniProtKB database, containing protein sequences from expertly curated genes, constituted a majority of each species' resolved genes and accounted for slightly less than one-third of the entire pan-gene pool (12,678 UniProtKB entities). Collectively, the three profile HMM peptide prediction databases identified a total of 10,351 unique gene entities, with Pfam predictably accounting for over half of all HMM hits (5,604 genes). Antibiotic resistance finders, like AMRFinder and BARRGD, identified a relatively small number of antimicrobial resistance genes, 0 – 4 on average. Similarly, ISfinder identified a relatively smaller quantity of insertion sequences in all species (ranging from 2 – 33). CD-hit identified 9,318 putative genes, for which no prior information was available.

We delineated genes based on their frequency to characterize the core, cloud, and shell genomes of the studied population(*40*), using classifications that follow the frequency cutoffs from Blaustein et al(*40*). The *S. enterica* cohorts had the most expansive cloud-genomes (73 % in the meat cohort, and 82 % clinical of total genes in cloud, Table 3). Conversely, the *S. sonnei* pan-genome was the most conserved, with 45% of its genes being core genes and only 52% of its genes belonging to the cloud genome (**Table S3**).

| Average (SD) | ***S. enterica,*** **meat tissue** | ***S. enterica,*** **clinical cohort** | ***E. coli*** | ***S. sonnei*** | ***V. parahaemolyticus*** | ***C. jejuni*** | ***C. coli*** |
|---|---|---|---|---|---|---|---|
| UniProtKB | 3,095 (53.9) | 3,080 (39.8) | 3,303 (50.0) | 3,154 (45.5) | 2,266 (108.0) | 886 (24.6) | 894 (25.0) |
| Pfam | 551 (44.4) | 540 (43.6) | 597 (25.6) | 495 (32.3) | 752 (28.6) | 283 (18.9) | 283 (19.4) |
| TIGRFAM | 82 (8.31) | 83 (11.4) | 80 (11.3) | 77 (11.5) | 58 (7.21) | 37 (4.46) | 35 (5.89) |
| HAMAP | 66 (6.16) | 64 (6.74) | 69 (4.15) | 59 (3.98) | 136 (4.02) | 115 (3.69) | 116 (6.75) |
| ISfinder | 18 (6.52) | 14 (4.92) | 33 (3.13) | 30 (4.37) | 10 (3.77) | 1 (0.84) | 2 (0.99) |
| AMRFinder | 0.2 (0.41) | 0.0 (0.12) | 3.7 (0.59) | 0.9 (0.27) | 0.0 | 0.0 | 0.0 |
| BARRGD | 1.0 (0.33) | 0.9 (0.38) | 0.0 (0.04) | 0.0 | 1.0 (0.0) | 0.4 (0.49) | 0.5 (0.50) |
| CD-hit | 114 (23.4) | 102 (18.6) | 150 (16.8) | 89 (16.6) | 247 (31.4) | 130 (20.6) | 138 (23.2) |
| **Total** | 3,927 (116) | 3,884 (93.9) | 4,236 (99.3) | 3,905 (101) | 3,470 (156) | 1,452 (54.5) | 1,469 (68.6) |

**Table S2**: **Major species group gene annotation statistics**. The average gene counts and associated standard deviations (SD) of each database or software platform are shown.

| **Gene type** | ***S. enterica,*** **meat tissue** | ***S. enterica,*** **clinical cohort** | ***E. coli*** | ***S. sonnei*** | ***V. parahaemolyticus*** | ***C. jejuni*** | ***C. coli*** |
|---|---|---|---|---|---|---|---|
| Core | 3,360 | 3,319 | 3,955 | 3,642 | 3,042 | 1,131 | 1,108 |
| Shell | 1,343 | 1,240 | 466 | 528 | 862 | 702 | 630 |
| Cloud | 12,585 | 19,641 | 6,887 | 3,930 | 4,158 | 6,887 | 2,692 |

**Table S3: Core-gene, shell-gene, and cloud gene counts for each major species group.**

223

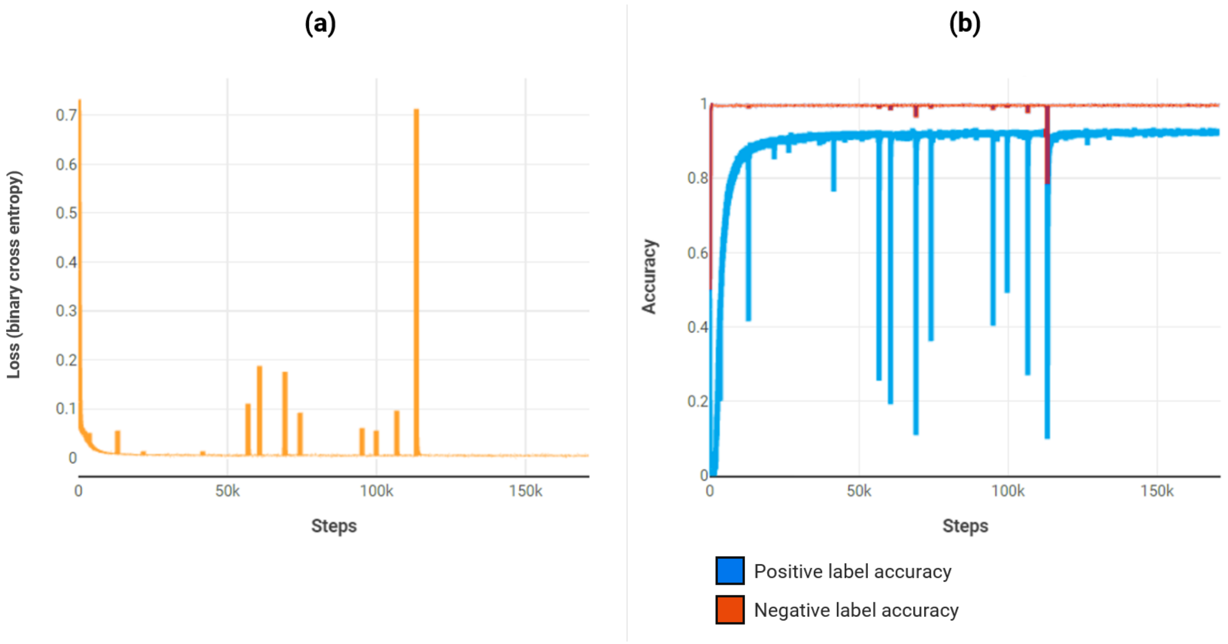


**Fig. S2: Training statistics over the course of on *S. enterica* as a single-species model**. **a** shows the reduction in binary cross entropy loss when training the Transformer to predict the AB genespace for single species *S. enterica* (meat tissue). **b** displays the corresponding training accuracy as the proportion of labels predicted correctly. The training accuracy is split between positive labels (blue) and negative labels (red).
